# Hypothalamic supramammillary neurons that project to the medial septum control wakefulness

**DOI:** 10.1101/2022.12.20.521335

**Authors:** Mengru Liang, Tingliang Jian, Wenjun Jin, Qianwei Chen, Xinyu Yang, Rui Wang, Jingyu Xiao, Zhiqi Yang, Xiang Liao, Xiaowei Chen, Liecheng Wang, Han Qin

**Affiliations:** Department of Anatomy, School of Basic Medical Sciences, Anhui Medical University, Hefei 230032, China; Brain Research Center and State Key Laboratory of Trauma, Burns, and Combined Injury, Third Military Medical University, Chongqing 400038, China; Farm Animal Genetic Resources Exploration and Innovation Key Laboratory of Sichuan Province, Sichuan Agricultural University, P. R. China; Center for Neurointelligence, School of Medicine, Chongqing University, Chongqing 400044, China; Chongqing Institute for Brain and Intelligence, Guangyang Bay Laboratory, Chongqing 400064, China

**Author notes:** Correspondence (X.C.), (L.W.), (H.Q.). These authors contributed equally (Mengru Liang, Tingliang Jian).

## Abstract

The hypothalamic supramammillary nucleus (SuM) plays a key role in controlling wakefulness, but the downstream target regions participating in this control process remain unknown. Here, using circuit-specific fiber photometry and single-neuron electrophysiology together with electroencephalogram, electromyogram and behavioral recordings, we find approximately half of SuM neurons that project to the medial septum (MS) are wake-active. Optogenetic stimulation of axonal terminals of SuM-MS projection induces a rapid and reliable transition to wakefulness from NREM or REM sleep, and chemogenetic activation of SuM^MS^ projecting neurons significantly increases wakefulness time and prolongs latency to sleep. Consistently, chemogenetically inhibiting these neurons significantly reduces wakefulness time and latency to sleep. Therefore, these results identify the MS as a functional downstream target of SuM and provide evidence for a causal role for this hypothalamic-septal projection in wakefulness control.

## Introduction

The supramammillary nucleus (SuM) is a hypothalamic region lying above the mammillary body, and provides abundant projections to numerous brain regions like the hippocampus, septum, frontal cortex, and cingulate cortex (Pan & McNaughton, 2004; Vertes, 1992). Recent advances in high-performance recording and manipulation techniques have enabled extensive studies of SuM functions, subsequently revealing its involvement in numerous processes such as episodic memory (Li et al., 2020; Qin et al., 2022), novelty detection (Chen et al., 2020), theta rhythm (Billwiller et al., 2020; Kocsis & Vertes, 1994), locomotion (Farrell et al., 2021), hippocampal neurogenesis (Li et al., 2022), and wakefulness (Pedersen et al., 2017). In particular, one previous study demonstrated that SuM glutamatergic neurons serve as a key node for arousal, and chemogenetic activation of SuM glutamatergic neurons, but not GABAergic neurons, produces sustained arousal (Pedersen et al., 2017). However, which downstream brain regions are involved in the SuM control of arousal remains unknown.

The medial septum (MS), which primarily contains cholinergic, GABAergic and glutamatergic neurons (Hajszan et al., 2004; Kiss et al., 1990), has been suggested to mediate different brain functions like locomotion (Fuhrmann et al., 2015), learning and memory (Boyce et al., 2016; Lecourtier et al., 2011), hippocampal theta generation (Buzsáki, 2002), and wakefulness (An et al., 2021; Osborne, 1994). Among these functions, MS glutamatergic neurons were shown to control wakefulness by activating lateral hypothalamic glutamatergic neurons (An et al., 2021). Furthermore, a recent study has demonstrated that SuM glutamatergic neurons project to MS glutamatergic neurons and are responsible for modulating the motivation for environmental interaction (Kesner et al., 2021). Based on this established anatomical connection and combined findings, we hypothesized that a SuM-MS projection may control wakefulness.

To test this hypothesis, we performed circuit-specific optical Ca^2+^ and optrode recordings in SuM-MS projection across sleep-wakefulness cycles. We identified a set of wake-active neurons in SuM that project to MS. Optogenetic or chemogenetic activation of SuM-MS projection induced behavioral and EEG arousal, and chemogenetic inhibition of this projection decreased wakefulness. Overall, our results reveal a critical role of the hypothalamic-septal projection for wakefulness control.

## Results

### SuM^MS^ projection terminals are strongly active during both wakefulness and REM sleep

SuM neurons have been reported to project to MS region (Vertes, 1988, 1992). We labeled the SuM-MS projection by local injection of an adeno-associated viral (AAV) vector to deliver the enhanced green fluorescent protein (eGFP) gene into SuM (Figure1-figure supplement 1A). Four weeks after injection, robust eGFP expression was observed in cell bodies within SuM (Figure1-figure supplement 1B), and the axonal terminals in MS were also labeled with eGFP (Figure1-figure supplement 1C). To further investigate the SuM to MS connection, retrograde AAV vector expressing eGFP was injected into MS (Figure1-figure supplement 1D). We verified that the expression area of eGFP was limited in MS (Figure1-figure supplement 1E) and the corresponding eGFP-labeled cell bodies were observed in SuM (f Figure1-figure supplement 1F).

Although both SuM and MS neurons have been shown to function as essential components in wakefulness (An et al., 2021; Pedersen et al., 2017), the activity of SuM neurons projecting to MS (SuM^MS^ projecting neurons) has not been recorded during sleep-wakefulness cycles. First, a circuit-specific fiber photometry system (Gunaydin et al., 2014; Qin et al., 2018; Qin et al., 2019) was used in conjunction with electroencephalogram (EEG) and electromyogram (EMG) recordings to observe Ca^2+^ activities at axonal terminals of SuM-MS projection in freely moving mice. For this purpose, AAV-syn-axon-jGCaMP7b (Broussard et al., 2018; Dana et al., 2019) was locally injected into SuM to express the Ca^2+^ indicator, jGCaMP7b, in axons of SuM neurons (Figure 1A). One month following virus injection, an optical fiber was implanted with the tip above MS to record activity at axonal terminals of SuM neurons and EEG-EMG electrodes were attached to the mouse cortical surface and neck muscles respectively, to define sleep-wakefulness states (Figure 1A). Virus expression and fiber tip location were verified by post-hoc histology after recording finished (Figure 1B). Notably, axonal terminals of SuM^MS^ projecting neurons had higher levels of Ca^2+^ activity during both wakefulness and REM sleep than that during NREM sleep (Figure 1C, also see statistics in Table Appendix 1 for all figures). In addition, these activities increased strongly during NREM-wakefulness and NREM-REM transitions, but sharply decreased during wakefulness-NREM transitions (Figure 1E-H). Taken together, these results suggested that SuM projects to MS and the Ca^2+^ activity in this projection is highly active during wakefulness and REM sleep.

**Figure 1.**
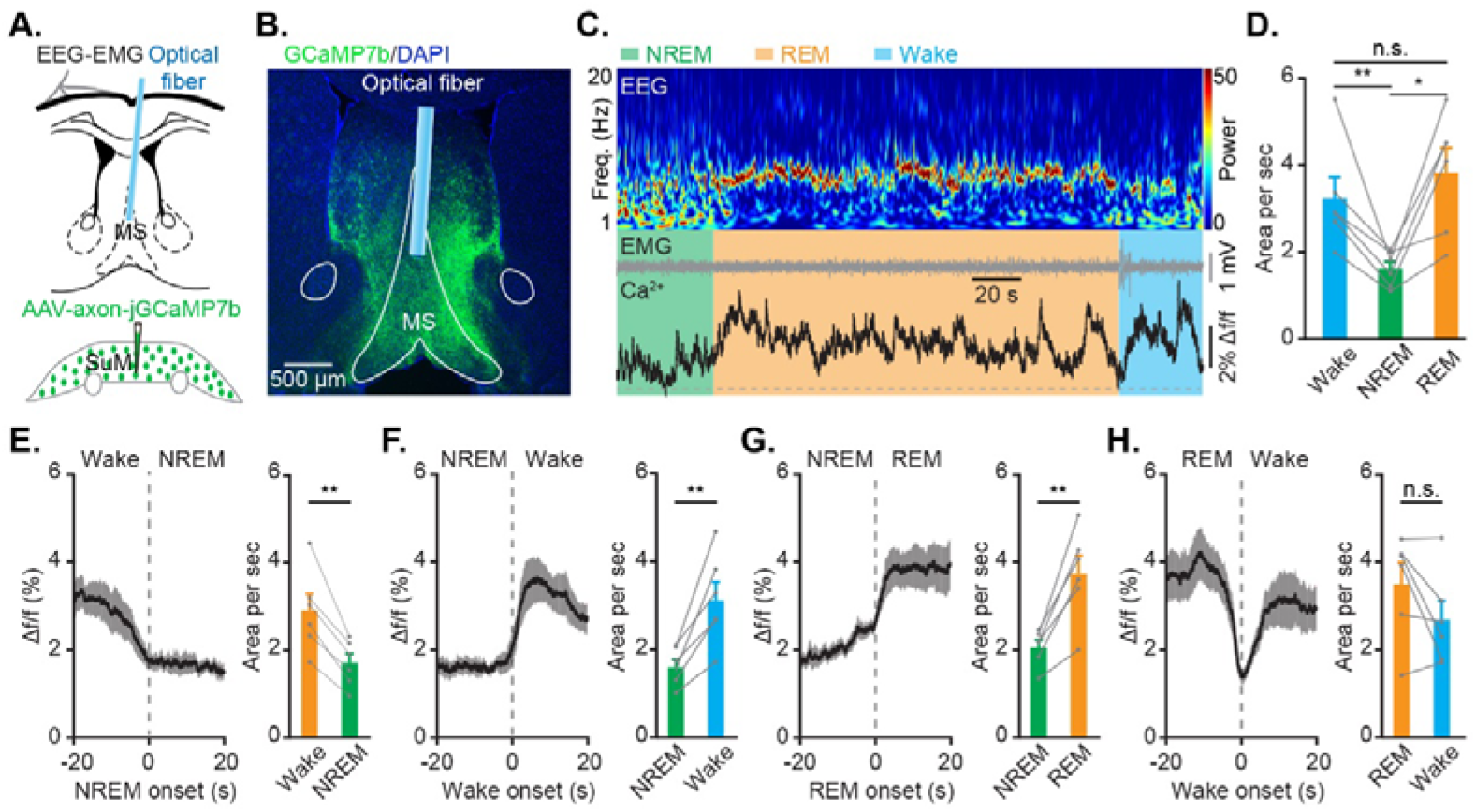
Strong activation of SuM-MS projection terminals during wakefulness and REM sleep. (**A**) Experimental design for virus injection into SuM, fiber implantation in MS, and EEG-EMG recording. (**B**) Post-hoc histological images showing the expression of jGCaMP7b at axonal terminals of SuM neurons and optical fiber location in MS. (**C**) Ca^2+^ activities in axonal terminals of SuM-MS projection across sleep-wakefulness cycles. Heatmap of EEG power spectrum (μV^2^). Freq., frequency. (**D**) Summary of the area under the curve per second during wakefulness, NREM sleep, and REM sleep. *n* = 6 mice, RMs 1-way ANOVA with LSD post-hoc comparison, \**p* < 0.05, \*\**p* < 0.01. (**E-H**) Ca^2+^ activities during brain state transitions: wakefulness-NREM (**E**), NREM-wakefulness (**F**), NREM-REM (**G**), and REM-wakefulness (**H**). *n* = 6 mice, paired *t* test, \*\**p* < 0.01.

### Identification of wake-active SuM^MS^ projecting neurons

To characterize the firing rates of SuM^MS^ projecting neurons at the single-cell level, we conducted optrode recordings across sleep-wakefulness cycles (Liu et al., 2020; Stark et al., 2012). Channelrhodopsin-2 (ChR2) was expressed specifically in SuM^MS^ projecting neurons by injecting a Cre-dependent retrograde AAV (retroAAV-Cre) into MS and, concurrently, an AAV vector carrying DIO-ChR2-mCherry into SuM. We then implanted an optrode in SuM to identify SuM^MS^ projecting neurons and record single-neuron activities (Figure 2A; see optrode locations in Figure2-figure supplement 1). A series of blue light pulses (450 nm, 2 Hz, 10 mW, 10 ms duration) were delivered to stimulate ChR2-expressing neurons. SuM neurons were then identified as MS projecting neurons based on light-induced spike which responses at short latency, low jitter, high success rate, and high correlation with spontaneous spike waveform (latency 3.6 ± 0.3 ms, jitter 0.8 ± 0.1 ms, success rate 95% ± 2%, correlation coefficient 0.92 ± 0.02, n = 23 neurons, Figure 2B-E).

**Figure 2.**
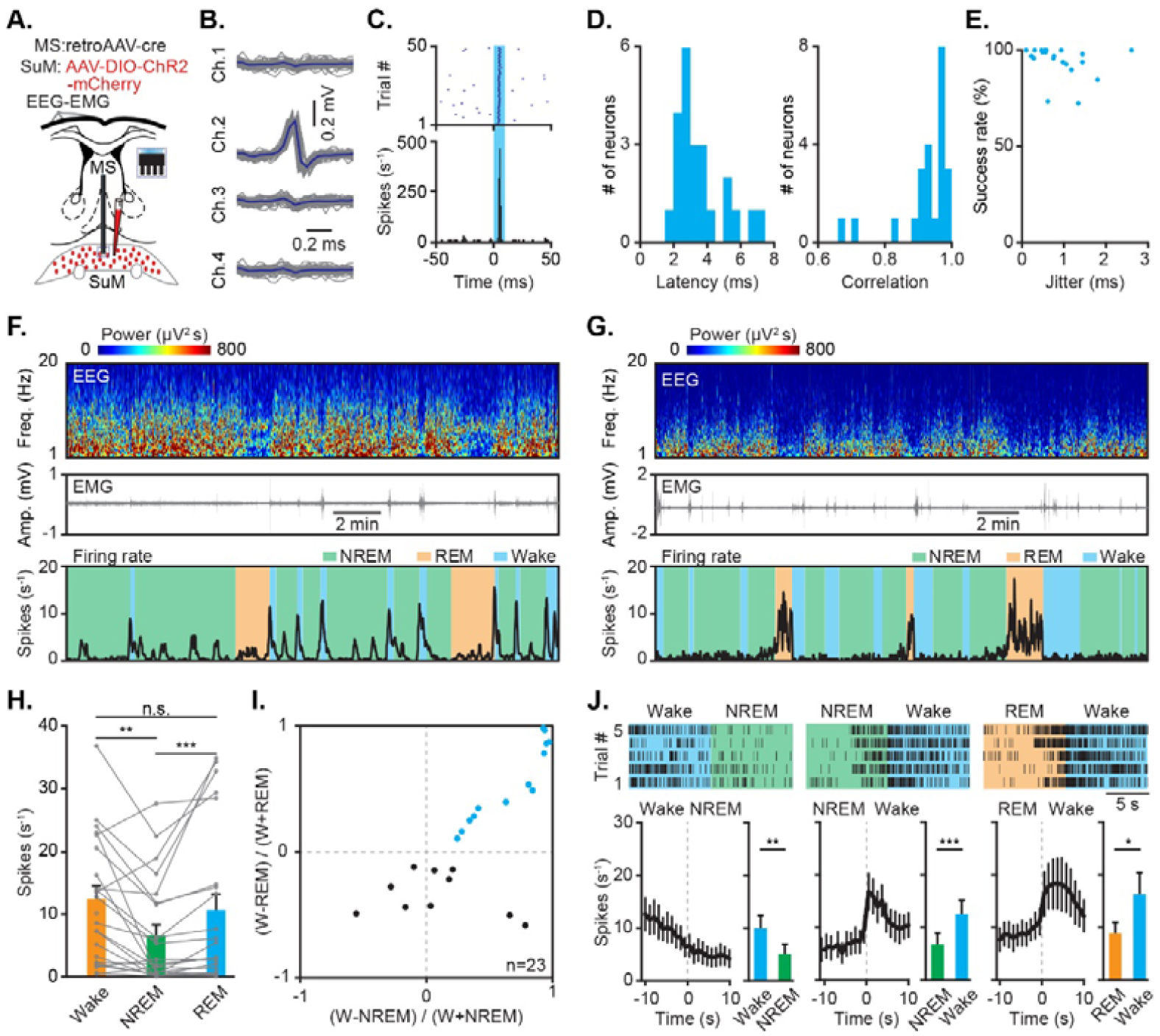
Optrode recording of wake-active SuM^MS^ projecting neurons. (**A**) Experimental design for retrograde labeling of SuM^MS^ projecting neurons, optrode recording in SuM, and EEG-EMG recording. (**B**) Waveforms of average light-invoked (blue) and individual spontaneous (gray) spikes from a representative SuM^MS^ projecting neuron. (**C**) Stimulus time histogram of neuronal spikes in (**B**). (**D**) Distributions of latencies before the first light-induced spikes (left), and correlation coefficients between light-induced spikes and spontaneous spikes (right) for all recorded SuM^MS^ projecting neurons. (**E**) Success rate versus temporal jitter of the first light-induced spikes for all recorded SuM^MS^ projecting neurons. (**F**) Firing rates of a representative wake-active neuron across sleep-wakefulness cycles. (**G**) Firing rates of a representative REM-active neuron across sleep-wakefulness cycles. (**H**) Summary of firing rates from 23 recorded SuM^MS^ projecting neurons (from 8 mice) in different states. Friedman’s ANOVA test and Wilcoxon signed-rank tests, \*\**p* < 0.01, \*\*\**p* < 0.001. (**I**) Firing rate modulation of SuM^MS^ projecting neurons, *n* = 23 neurons from 8 mice. (**J**) Firing rate of wake-active SuM^MS^ projecting neurons during state transitions: wakefulness-NREM, left; NREM-wakefulness, middle; REM-wakefulness, right. Top: example of a wake-active SuM^MS^ projecting neuron during five trials of different state transitions. Bottom left: average firing rates during state transitions. Bottom right: summary of firing rates of 10 s before and after state transitions, paired *t* test, \**p* < 0.05, \*\**p* < 0.01, \*\*\**p* < 0.001, *n* = 13 neurons.

We found that two groups of neurons showed distinct firing features across sleep-wakefulness cycles. Neurons in one group significantly increased firing rates following the transition from NREM or REM sleep to wakefulness, and significantly decreased their firing rates following the switch from wakefulness to NREM sleep (wake-active neuron, see example in Figure 2F). Neurons in the other group significantly increased firing rates following the transition from NREM sleep to REM sleep and significantly decreased their firing rates following the switch from REM sleep to wakefulness (REM-active neuron, see example in Figure 2G). We summarized the firing rates of all SuM^MS^ projecting neurons in wakefulness, NREM sleep, and REM sleep states. Consistent with the results of Ca^2+^ activity at axonal terminals, the firing rates of SuM^MS^ projecting neurons were significantly higher during wakefulness and REM sleep than that during NREM sleep (Figure 2H; wakefulness, 12.4 ± 2.1 Hz; REM, 10.6 ± 2.6 Hz; NREM, 6.6 ± 1.7 Hz; Friedman’s ANOVA and Wilcoxon signed-rank tests, *n* = 23 neurons from 8 mice, wakefulness versus NREM, *P* = 0.001, REM versus NREM, *P* = 0.0002, wakefulness versus REM, *P* = 0.24).

Analysis of firing modulation by SuM^MS^ projecting neurons during these three states followed by calculation of firing rates in wake-active neurons during state transitions (Figure 2I) revealed that these wake-active neurons had higher firing rates during NREM-wakefulness or REM-wakefulness transitions, but lower firing rates during wakefulness-NREM transitions (Figure 2J). These results established that wake-active neurons were indeed present in SuM-MS projection, likely contributing to control of wakefulness.

### Stimulating SuM-MS projection promotes wakefulness

To determine whether the SuM-MS projection indeed play a causal and key role in the control of wakefulness, optogenetic activation was applied in MS to activate ChR2-expressing axonal terminals of SuM neurons (Figure 3A). To this end, we injected AAV-ChR2-mCherry into SuM to express ChR2 in axonal terminals of SuM-MS projection (see virus expression in Figure 3B and C) and an optical fiber was subsequently implanted into MS to deliver blue light. EEG-EMG electrodes were attached to monitor activity during sleep-wakefulness cycles, and axonal terminals of SuM-MS projection were optogenetically activated for 20 s (473 nm, 15 mW, 10 ms duration) after the onset of stable NREM or REM sleep. The activation induced transition from NREM sleep to wakefulness in a frequency-dependent manner (example in Figure 3D; statistics in Figure 3E and F). The success rate of transition from NREM sleep to wakefulness after 20-Hz optogenetic activation was 100%, with a latency to wakefulness of 2.0 ± 0.3 s (SuM-DG ChR2 control: 65.8 ± 11.8 s, SuM-MS mCherry control: 65.1 ± 6.0 s). Transition to wakefulness was also induced upon 20-Hz optogenetic activation of this projection during REM sleep (example in Figure 3G; statistics in Figure 3H and I), with an 89% ± 8% success rate and a latency to wakefulness of 16.4 ± 6.7 s (SuM-DG ChR2 control: 64.7 ± 3.7 s, SuM-MS mCherry control: 59.2 ± 7.5 s).

**Figure 3.**
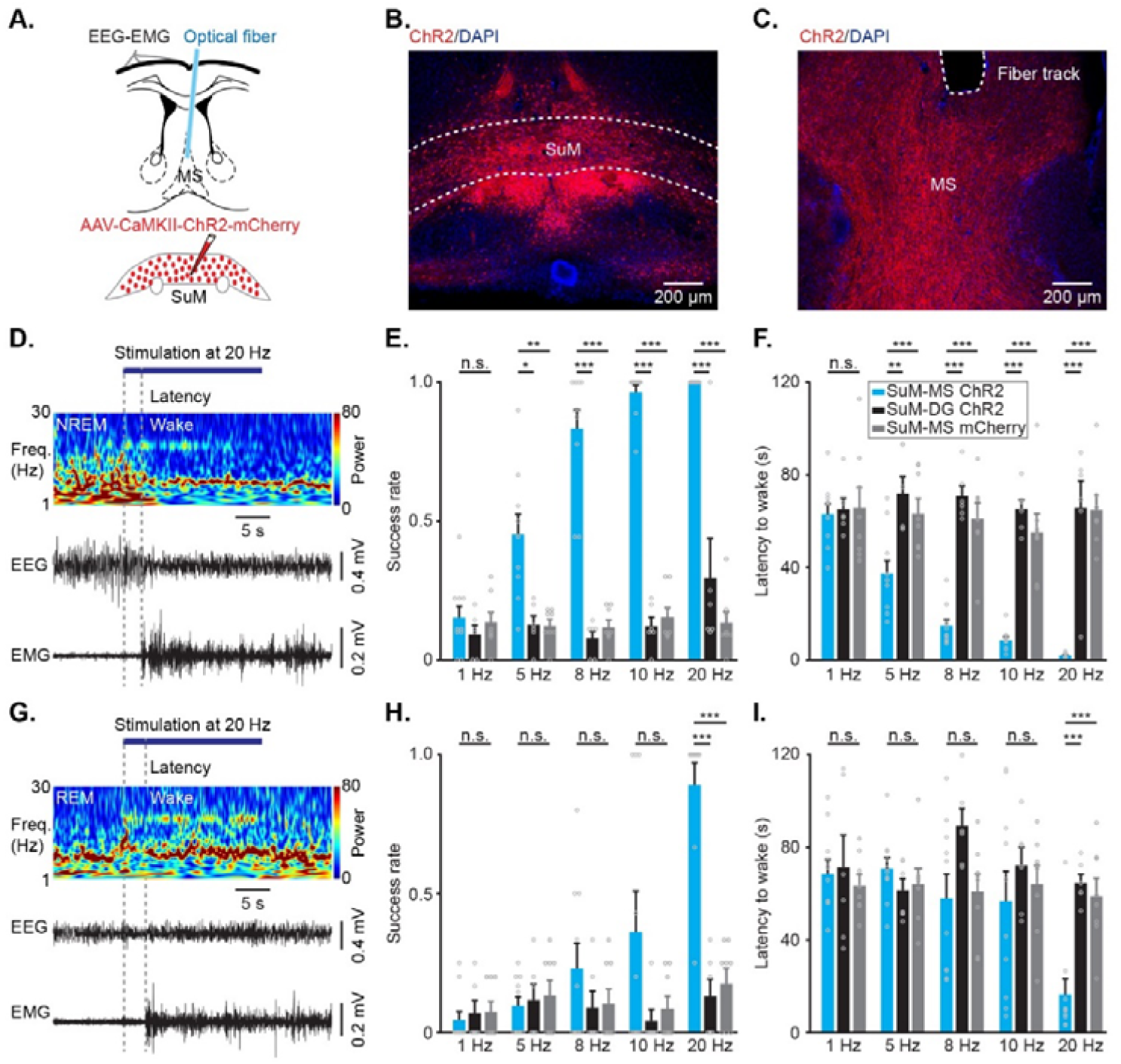
Optogenetic stimulation at axonal terminals of SuM neurons in MS induces wakefulness. (**A**) Experimental design for virus injection into SuM, fiber implantation in MS, and EEG-EMG recording. (**B-C**) Representative images showing mCherry-labelled somas in SuM (**B**), and mCherry-labelled axonal terminals of SuM neurons and the track of fiber in MS (**C**); Scale bar = 200 μm. (**D**) Representative EEG power spectrum (μV^2^) and EEG-EMG trace data around 20-Hz stimulation during NREM sleep. Freq., frequency. (**E**) Summary of the success rate for inducing wakefulness from NREM sleep in different groups; SuM-MS ChR2, *n* = 10, SuM-DG ChR2, *n* = 6, SuM-MS mCherry, *n* = 8; RMs 2-way ANOVA with Sidak post-hoc comparison test, \**p* < 0.05, \*\**p* < 0.01, \*\*\**p* < 0.001. (**F**) Summary of the latency to wakefulness from NREM sleep; RMs 2-way ANOVA with Sidak post-hoc comparison test, \*\**p* < 0.01, \*\*\**p* < 0.001. (**G**) Representative EEG power spectrum (μV^2^) and EEG-EMG trace data around 20-Hz stimulation during REM sleep. (**H**) Summary of the success rate for inducing wakefulness from REM sleep; SuM-MS ChR2, *n* = 10, SuM-DG ChR2, *n* = 6, SuM-MS mCherry, *n* = 8; RMs 2-way ANOVA with Sidak post-hoc comparison test, \*\*\**p* < 0.001. (**I**) Summary of the latency to wakefulness from REM sleep; RMs 2-way ANOVA with Sidak post-hoc comparison test, \*\*\**p* < 0.001.

To verify the above results, SuM^MS^ projecting neurons were selectively chemogenetically activated by specific labeling with an engineered G_i_-coupled hM3Dq receptor (Armbruster et al., 2007). We injected retroAAV-Cre into MS and AAV-DIO-hM3Dq-mCherry into SuM (Figure 4A) to label these SuM^MS^ projecting neurons. And robust expression of hM3Dq-mCherry in SuM neurons was confirmed by post-hoc histological analysis (Figure 4B). Immunostaining for c-Fos protein (a marker of active neurons) (Adamsky et al., 2018; Zhou et al., 2018) showed that SuM^MS^ projecting neurons were activated after application of the synthetic ligand clozapine-N-oxide (CNO, 1 mg/Kg). In hM3Dq-positive neurons, c-Fos expression was significantly higher in the CNO application group than in saline-treated control animals (Figure 4B and C). At behavioral level, intraperitoneal injection with CNO at the start of the light period resulted in significantly greater wakefulness (Figure 4D-F; RMs 2-way ANOVA test, *P* = 0.0008, F = 17.9, *n* = 8 mice) and latency to first sleep was significantly longer in the CNO group (7.1 ± 0.5 h vs 0.7 ± 0.1 h, Paired t test, *P* < 0.001, *n* = 8 mice) compared with that in the saline control (Figure 4G). These experiments thus demonstrated that optogenetic and chemogenetic activation of SuM^MS^ projecting neurons could promote wakefulness and maintain at this state.

**Figure 4.**
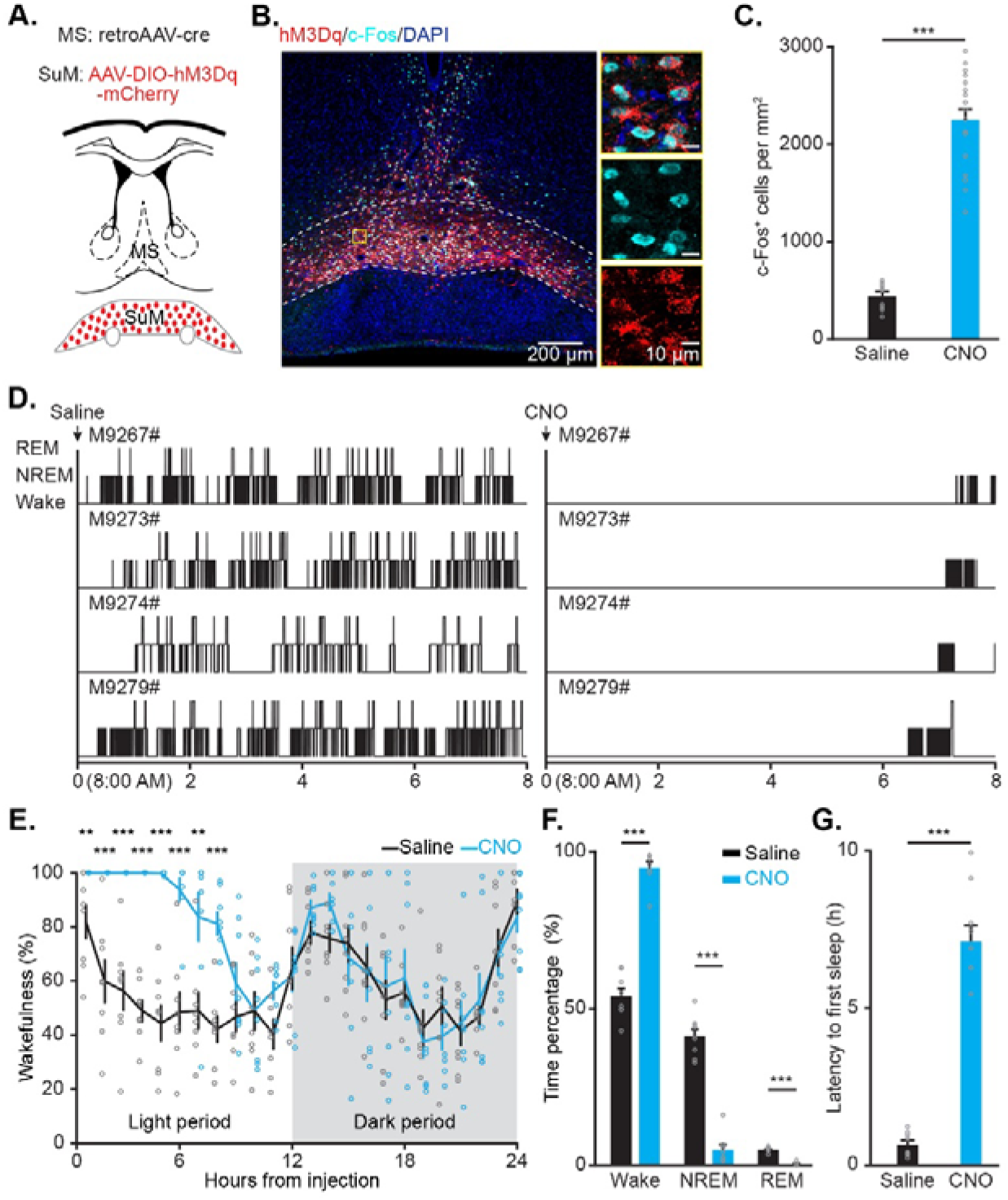
Chemogenetic activation of SuM^MS^ projecting neurons increases wakefulness. (**A**) Schematic for virus injection of retroAAV-Cre into MS and AAV-DIO-hM3Dq-mCherry into SuM. (**B**) Post-hoc histological image of hM3Dq expression in SuM and c-Fos expression after intraperitoneal injection of CNO; yellow rectangle indicates magnified area. (**C**) Summary of c-Fos^+^ neurons in saline- and CNO-treated mice. Saline: *n* = 9 brain sections from 3 mice, CNO: *n* = 18 brain sections from 6 mice; Wilcoxon rank-sum test, \*\*\**p* < 0.001. (**D**) Hypnograms of hM3Dq-mCherry mice in the 8 h following saline (left) or CNO (right) injection. (**E**) Hourly percentage of time in wakefulness across 24-hour recording period after saline or CNO injection in hM3Dq-mCherry mice, *n* = 8 mice, RMs 2-way ANOVA with Sidak post-hoc comparison test. (**F**) Percentage of time in each state during the first 8 h after saline or CNO injection, *n* = 8 mice; Wilcoxon rank-sum test, \*\*\**p* < 0.001. (**G**) Summary of the latency to first sleep after saline or CNO injection, *n* = 8 mice; paired *t* test, \*\*\**p* < 0.001.

### Chemogenetic inhibition of SuM^MS^ projecting neurons reduces wakefulness

To further examine how chemogenetic inhibition of SuM^MS^ projecting neurons affects control of wakefulness, expression of an engineered G_i_-coupled hM4Di receptor in SuM^MS^ projecting neurons was induced by injecting retroAAV-Cre into MS and AAV-DIO-hM4Di-mCherry into SuM (Figure 5A). Immunostaining detection of c-Fos verified that CNO injection led to the inhibition of SuM^MS^ projecting neurons in hM4Di-expressing mice, indicated by lower c-Fos signal in the CNO group than in saline control (Figure 5B-C). Examination of behavioral states (Figure 5D-E) indicated that mice in the CNO group had shorter latency to first sleep than that in saline control (14.5 ± 1.8 vs 28.8 ± 3.3 min, Paired t test, *P* = 0.002, *n* = 10 mice) and wakefulness was significantly reduced during the first 2 h in the CNO-treated mice (Figure 5D and F). These results thus indicated that acute inhibition of SuM^MS^ projecting neurons reduces wakefulness and increases sleep.

**Figure 5.**
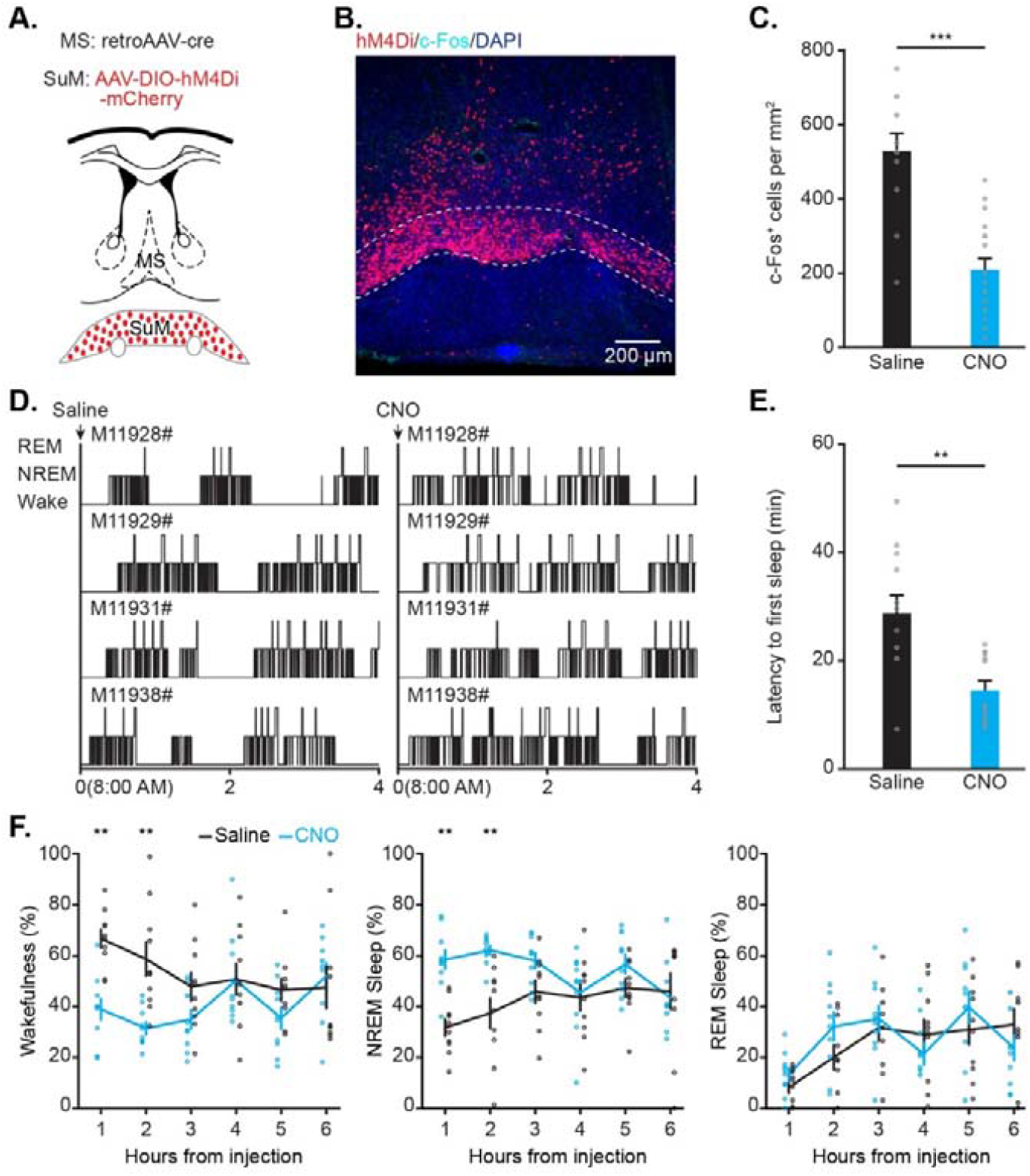
Chemogenetic inhibition of SuM^MS^ projecting neurons reduces wakefulness. (**A**) Schematic for injection of retroAAV-cre into MS and AAV-DIO-hM4Di-mCherry into SuM. (**B**) Post-hoc histological image of hM4Di expression in SuM and c-Fos expression after intraperitoneal injection of CNO. (**C**) Summary of c-Fos^+^ neurons in saline- and CNO-treated mice. Saline: *n* = 12 brain sections from 3 mice, CNO: *n* = 18 brain sections from 5 mice; unpaired *t* test, \*\*\**p* < 0.001. (**D**) Hypnograms of hM4Di-mCherry mice in the 4 h following saline (left) or CNO (right) injection. (**E**) Summary of the latency to first sleep after saline or CNO injection, *n* = 10 mice; paired *t* test, \*\**p* < 0.01. (**F**) Hourly percentage of time in wakefulness (left), NREM sleep (middle), and REM sleep (right) during the 6 h following saline or CNO injection in hM4Di-mCherry mice; *n* = 10 mice; RMs 2-way ANOVA with Sidak post-hoc comparison test, \*\**p* < 0.01.

## Discussion

Control of wakefulness requires multiple brain regions that span across the entire neural networks (Brown et al., 2012; Liu & Dan, 2019; Saper & Fuller, 2017). And identification of wake-active neurons is a necessary step in resolving the mechanism(s) underlying the regulation of wakefulness. For example, monoaminergic neurons in the ascending activating system (Gu et al., 2022; Kayama & Koyama, 2003; Liu & Dan, 2019; Scammell et al., 2017) and orexin neurons in lateral hypothalamus (Lee et al., 2005; Mileykovskiy et al., 2005) have been identified as wake-active neurons that are essential for wakefulness control. Here, using optical fiber and optrode recordings, we found a group of wake-active MS projecting neurons in SuM (Figures 1 and 2). Optogenetic stimulation of axonal terminals from these SuM^MS^ projecting neurons was sufficient to induce a rapid and reliable transition to wakefulness from sleep (Figure 3). Retrograde projection-specific labeling combined with chemogenetic manipulation revealed that the SuM^MS^ neurons play an essential role in maintaining wakefulness (Figures 4 and 5).

Previous work has shown that MS glutamatergic neurons are all wake-active and involved in wakefulness control (An et al., 2021), and SuM neurons can activate hippocampal neurons during REM sleep and locomotion (Farrell et al., 2021; Li et al., 2022; Renouard et al., 2015). In addition, our previous study revealed a REM-active pattern in all SuM-hippocampus projecting neurons and that these neurons are critical for episodic memory consolidation (Qin et al., 2022). However, for SuM^MS^ projecting neurons, the firing patterns appear more complicated in different behavioral states. SuM neurons exhibit high activity during exploration and approach behaviors, but low activity during sucrose consumption (Kesner et al., 2021). Our results here show that approximately half (13/23) of the SuM^MS^ projecting neurons are wake-active, while about 39% (9/23) neurons are REM-active. It is possible that these REM-active SuM^MS^ projecting neurons might participate in certain REM sleep-related functions, such as memory consolidation (Boyce et al., 2016; Kumar et al., 2020; Qin et al., 2022) or cortical plasticity (Peever & Fuller, 2017; Sterpenich et al., 2014).

SuM mainly contains Vgat, Vglut2, Tac1 and Nos1 neurons (Farrell et al., 2021; Kesner et al., 2021; Pan & McNaughton, 2004; Pedersen et al., 2017), among which Tac1 neurons project to the septum, hippocampus and other regions, functioning in the control of locomotion (Farrell et al., 2021). By contrast, SuM Vglut2 neurons promote arousal, while Nos1/Vglut2 neurons together contribute to theta rhythm in the hippocampus during REM sleep (Pedersen et al., 2017). In particular, SuM^MS^ projecting neurons are well-established to be primarily glutamatergic (Kesner et al., 2021), and SuM Vglut2 neurons can monosynaptically innervate MS Vglut2 neurons by releasing glutamate (Kesner et al., 2021). Thus, wake-active SuM^MS^ projecting neurons identified here are likely to be glutamatergic and innervate MS Vglut2 neurons, which are known to control wakefulness by activating lateral hypothalamus glutamatergic neurons (An et al., 2021; Wang et al., 2021).

## Materials and methods

### Animals

12-20-week-old C57BL/6J mice (male) were used in the recording and manipulation experiments. Mice were housed in groups under a constant temperature (21-24°C) and humidity (50%-60%), while those implanted with optical fibers or optrodes were maintained in individual cages. All mice were housed under a 12/12-hour light/dark cycle (with lights on at 7:00 am), and had free access to food and water. All experimental procedures were conducted according to the protocols and guidelines of the Third Military Medical University Animal Care and Use Committee.

### Virus

AAV2/8-EF1α-eGFP (titer: 1.49×10^13^ viral particles/mL) and retroAAV2/2 Plus-EF1α-eGFP (titer: 1.92 × 10^13^ viral particles/mL) were used for tracing experiments. AAV2/9-Syn-axon-jGCaMP7b (titer: 2.17 × 10^13^ viral particles/mL) was used for Ca^2+^ recording. AAV2/9-EF1α-DIO-hChR2-mCherry (titer: 3.67×10^13^ viral particles/mL) and retroAAV2/2 Plus-Syn-Cre (titer: 1.92 × 10^13^ viral particles/mL) were used for optrode recording. AAV2/9-CaMKII-hChR2-mCherry (titer: 1.72 × 10^13^ viral particles/mL), AAV2/9-CaMKII-mCherry (titer: 1.72 × 10^13^ viral particles/mL), retroAAV2/2 Plus-Syn-Cre (titer: 1.92 × 10^13^ viral particles/mL), AAV2/9-DIO-hM3Dq-mCherry (titer: 1.00 × 10^12^ viral particles/mL), AAV2/9-DIO-hM4Di-mCherry (titer: 1.00 × 10^12^ viral particles/mL), and AAV2/9-DIO-mCherry (titer: 1.00 × 10^12^ viral particles/mL) were used for optogenetic and chemogenetic manipulations. All of the AAV constructs mentioned above were purchased from Taitool Bioscience Co., Ltd. (Shanghai, China) or Obio Biotechnology Co., Ltd. (Shanghai, China).

### Optrode construction for *in vivo* recording

An optrode consisted of an optical fiber (200 μm diameter, NA 0.37) and four custom-made tetrodes. A tetrode was grouped by four insulated tungsten wires (25 μm diameter, California Fine Wire). The four tetrodes were arranged into a line with a spacing of ∼200 μm and fixed by a fused silica capillary tube, and then mounted onto a micro-drive for vertical movement. The optical fiber was fixed to tetrodes with the tips being ∼500 μm shorter than the tetrode tips. The light from a laser diode (450 nm) was collimated to the optical fiber at the opposite end with a maximal light intensity measured by an optical power meter (PM100D, Thorlabs). Optical adhesive was used to connect the laser diode and optical fiber.

### Surgical procedures

For all surgeries, mice were anesthetized with 3% isoflurane in oxygen for 3-5 min and then placed into a stereotaxic frame with an isoflurane concentration maintained at 1%-2%. A heating pad was put under the mice to maintain a temperature of ∼37 °C throughout the surgery process. After surgery, the mice were placed back in warm cages and allowed to fully recover. Moreover, they received one dose of dexamethasone sodium phosphate (1mg/ml, 0.1ml/10g/d) and ceftriaxone sodium (50mg/ml, 0.1ml/10g/d) per day by intraperitoneal injection for 3 consecutive days to reduce inflammation (Li et al., 2018; Zhao et al., 2020).

For virus injection, 8-12-week-old mice were used. A glass pipette (tip diameter: 10-20 μm) was inserted through a small craniotomy (0.5 × 0.5 mm) to deliver the virus to specific brain areas. To express eGFP in the SuM-MS projection, ∼50 nL of AAV-eGFP was injected into SuM (AP: -2.8 mm, ML: 1.0 mm, 5° angle towards the midline, DV: 5.0 mm) or ∼200 nL of retro AAV2/2-eGFP was injected into MS (AP: 1.0 mm, ML: 0.5 mm, 5° angle towards the midline, DV: 3.8 mm). To express ChR2 or mCherry in the axons of SuM-MS projection, ∼200 nL of AAV-CaMKII-hChR2-mCherry or AAV-CaMKII-mCherry was injected into SuM. To express ChR2, hM3Dq, hM4Di, or mCherry specifically in SuM^MS^ projecting neurons, ∼200 nL of retroAAV2/2-Cre was injected into MS concurrent with ∼200 nL of AAV-DIO-ChR2-mCherry, AAV-DIO-hM3Dq-mCherry, AAV-DIO-hM4Di-mCherry or AAV-DIO-mCherry was injected into SuM. The viruses were allowed sufficient expression for about one month before subsequent experiments.

For fiber implantation, mice injected with AAV-axon-jGCaMP7b, AAV-CaMKII-ChR2-mCherry, or AAV-CaMKII-mCherry were used. To record Ca^2+^ activity, a self-made fiber probe (Qin et al., 2019) was prepared with an optical fiber (200 μm diameter, NA 0.53, MFP_200/230/900-0.53, Doric lenses) glued into a mental cannula (ID:0.51 mm, OD: 0.82 mm) after the end face was cut flat. To deliver blue light for optogenetic excitation, optical fiber ferrules (200 μm diameter, NA 0.37, Hangzhou Newdoon Technology Co., Ltd) were used. The prepared fiber probe was inserted through a small cranial window above MS (AP: 1.0 mm, ML: 0.5 mm, 5° angle towards the midline) to a depth of 3.5 mm. Blue light-curing dental cement (595989WW, Tetric EvoFlow) was applied to fix the probe to the skull. Further reinforcement was achieved with a common dental cement mixture in super glue.

For optrode implantation, mice expressing ChR2 in SuM^MS^ projecting neurons were used. Similarly, the previously described optrode was inserted after a craniotomy above SuM was made. The implantation depth was 4.7 mm from the dura. After a full recovery (the body weight started to increase), the optrode was gradually advanced to the target depth of ∼5.0 mm by micro-drive.

For EEG-EMG electrodes implantation, three EEG electrodes made by stainless steel screws were inserted into the craniotomy holes, with two above the frontal lobe (AP: 1.3 mm, ML: ± 1.2 mm) and the third one above the parietal lobe (AP: - 3.2 mm, ML: 3.0 mm). Two fine-wire EMG electrodes were inserted into the neck musculature for EMG recording.

Before all recording and manipulation experiments, mice were connected to optical and electrophysiological recording cables in the recording cages to habituate for 3 consecutive days.

### Fiber recording

A previously described fiber photometry system was used for Ca^2+^ recording (Qin et al., 2018; Qin et al., 2022). The recording was performed in jGCaMP7b-expressing mice with a fiber probe implanted in MS. Ca^2+^ activity (2 KHz), EEG-EMG signals (200 Hz), and behavioral videos (25 Hz) were simultaneously recorded across sleep-wakefulness cycles. Offline event makers were used to synchronize these three forms of signals.

### Optrode recording

Excitation light pulses (450 nm wavelength, 10 ms duration, ∼10 mW intensity, 0.5 s interval) were applied in optrode-implanting mice to identify SuM^MS^ projecting neurons. Units evoked by light stimulation with short spike latency (< 8 ms for all the units in our data) and high response reliabilities (> 73% for all the units in our data) were identified as ChR2-positive neurons. Then electrophysiological recordings (sampled at 20 KHz), EEG-EMG recording, and behavioral video recordings were simultaneously conducted across sleep-wakefulness cycles in the light phase. After all recordings were finished, an electrical lesion (current with 30 μA intensity and 12 s duration) was made to verify the recording sites.

### Optogenetic stimulation

473 nm blue laser light (MBL-III-473, Changchun New Industries) was delivered through an optical fiber ferrule under the control of a self-written program on the LabVIEW platform (LabVIEW 2014, National Instrument). The intensity of the light was measured with an optical power meter (PM100D, Thorlabs) and calibrated to ∼15 mW at the fiber tip. Stimulation pulses with 10 ms in duration at 1/5/8/10/20 Hz were delivered randomly during NREM or REM sleep. The EEG and EMG signals were manually monitored by experimenters in real time, and stimulation was applied after 20 s from the onset of stable NREM or REM sleep in the light phase.

### Chemogenetic manipulation

Chemogenetic manipulations were applied to hM3Dq or hM4Di-expressing mice after EEG-EMG electrodes implantation. After recovery, CNO (1 mg/kg, dissolved in saline, 0.3 mL) or an equal volume of saline was intraperitoneally injected. EEG-EMG signals and behavioral videos were recorded 2 h before drug applications and lasted for 24 hours.

### Sleep structure analysis

All EEG-EMG signals were first band-filtered (EEG: 0.5-30 Hz, EMG: 10 - 70 Hz). Then signals were divided into non-overlapping epochs of 4 s for analysis. The NREM sleep, REM sleep, or wakefulness state was automatically defined according to the amplitude of EMG and δ/θ power of the EEG spectrum by sleep analysis software (SleepSign for Animal, Kissei Comtec). NREM sleep was characterized by high amplitude in the EEG δ band (0.5-4 Hz) and low amplitude of EMG activity. REM sleep was characterized by low amplitude in the EEG δ band and high amplitude in EEG θ band (4-10 Hz), without tonic EMG activity. Wakefulness was characterized by high EMG activity and low amplitude of EEG activity. The automatically defined results were reviewed and manually corrected. The cumulative duration of NREM sleep, REM sleep, and wakefulness were summarized by a self-written MATLAB program.

### Histology

All mice used above were perfused with 4% paraformaldehyde (PFA) in PBS. The brains were sectioned into 50-μm slices after being dehydrated with 15% sucrose in 4% PFA for 24 h. Brain sections were imaged by a wide-field fluorescence microscope (Olympus, BX51) or confocal microscope (Zeiss, LSM 700) after being stained with DAPI. For immunohistochemistry, mice expressing hM3Dq or hM4Di were perfused 1.5 h after CNO or saline injection and sectioned as described above. Brain sections were blocked and incubated with primary antibodies (rabbit anti-c-Fos 1:200, ab190289, Abcam, RRID: AB_2737414), as previously described (Qin et al., 2020; Zhang et al., 2016). The number of c-Fos expressing neurons was manually counted by experiment-blinded analysts.

### Data analysis and statistics

The data of Ca^2+^ signals were analyzed as previously described (Qin et al., 2018; Qin et al., 2019). Briefly, all Ca^2+^ signals were filtered by Savitzky-Golay FIR smoothing filter with 50 side points and a 3rd-order polynomial. Then Ca^2+^ signals were calculated into Δf/f by the formula of Δf/f = (f - f_baseline_) / f_baseline_, where f_baseline_ represents the baseline fluorescence obtained during recording. To quantify the Ca^2+^ signals during sleep-wakefulness cycles, we identified the arousal state based on synchronous EEG-EMG signals. The area under Ca^2+^ signals was used for statistical analysis (Qin et al., 2022).

The raw extracellular electrophysiological data were analyzed as described previously (Qin et al., 2018; Qin et al., 2022). Events that exceeded an amplitude threshold of four standard deviations above the baseline were saved for subsequent spike sorting analysis. All detected events for each tetrode were sorted in the toolbox MClust based on the features of waveforms (Schmitzer-Torbert & Redish, 2004). The firing rates of each unit were calculated in a sliding time bin of 2 s (0.1 s interval). Units were classified according to the firing rates in NREM sleep, REM sleep, and wakefulness. We analyzed the spectral profiles of EEG activity by a self-designed MATLAB program (Ren et al., 2018). The EEG data were calculated by fast Fourier transformation with a frequency resolution of 0.15 Hz.

Statistical analyses were performed in MATLAB and SPSS22.0 (Table Appendix 1). Normality tests were analyzed between samples. Parametric tests (paired and unpaired *t*-tests, RMs 1-way ANOVA with LSD post hoc comparison, 1-way ANOVA with LSD post hoc comparison, and RMs 2-way ANOVA with Sidak’s post hoc comparison) were subsequently applied if normality or equal variance was achieved. Otherwise, non-parametric tests (Wilcoxon signed-rank test, Wilcoxon rank-sum test, Kruskal-Wallis test with Tukey post hoc comparison, and Friedman’s ANOVA test) were applied. All tests were two-tailed. All summary data were from individual mice and represented as mean ± SEM.

## Acknowledgements

The authors are grateful to Ms. Jia Lou for help in composing and layout editing of the figures. This work was supported by grants from the National Key R&D Program of China (2021YFA0805000), the National Natural Science Foundation of China (31925018, 32127801, 31921003, 81971236, 32200838), Chongqing Basic Research grants (cstc2019jcyjjqX0001) to X.C. and Fundamental Research Funds for the Central Universities (2022CDJXY-024) to H.Q.. X.C. is a member of the CAS Center for Excellence in Brain Science and Intelligence Technology.

## Author contribution

Conceptualization and methodology, H.Q., L.W., and X.C.; software programming, W.J. and X.L.; data curation, M.L., T.J., and H.Q.; investigation, M.L., T.J., Q.C., X.Y., and Z.Y.; technical support, R.W., and J.X.; writing-original draft, M.L.; writing – review & editing, H.Q. and X.C.; funding acquisition, L.W., X.C. and H.Q.; resources, M.L., T.J., W. J. and H.Q.; supervision, H.Q. and X.C. All authors read and commented on the manuscript.

## Supplementary files

Table Appendix 1. Statistics summary in the study, related to Figures 1–5. Only statistically significant (p < 0.05) results are reported.

## Data availability

All data needed to evaluate the conclusions in the paper are present in the manuscript and/or the figure supplement.

---

**Figure 1-figure supplement 1.**
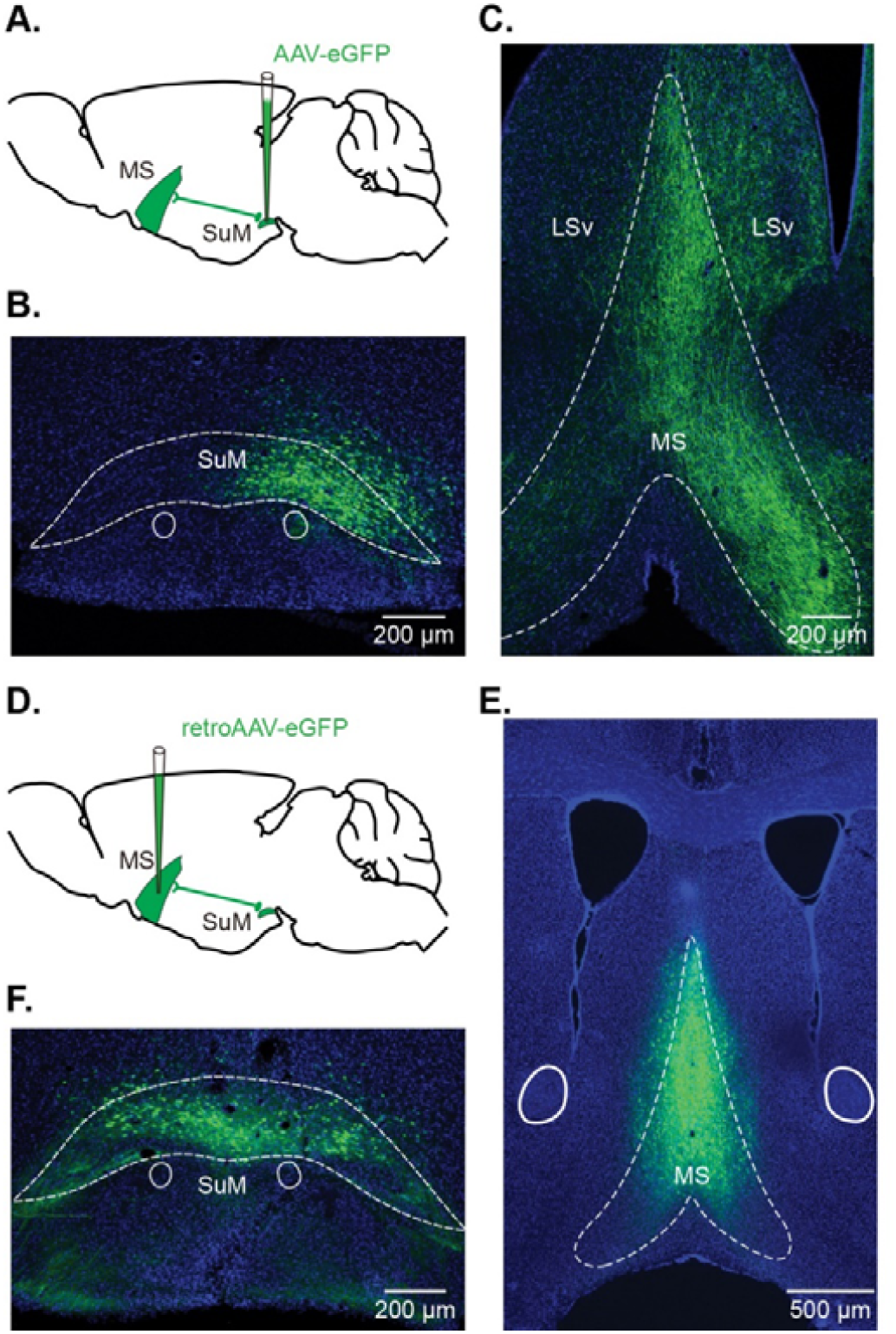
SuM neurons project to MS. (**A**) Diagram of AAV-eGFP injection into SuM. (**B-C**) Representative histological images of eGFP-labeled cell bodies in SuM (**B**) and their axon fibers in MS (**C**). The same experiments were performed in 5 mice. (**D**) Diagram of retroAAV-eGFP injection into MS. (**E-F**) Representative histological images of injection area in MS (**E**) and eGFP-labeled cell bodies in SuM (**F**). The same experiments were performed in 6 mice.

**Figure 2-figure supplement 1.**
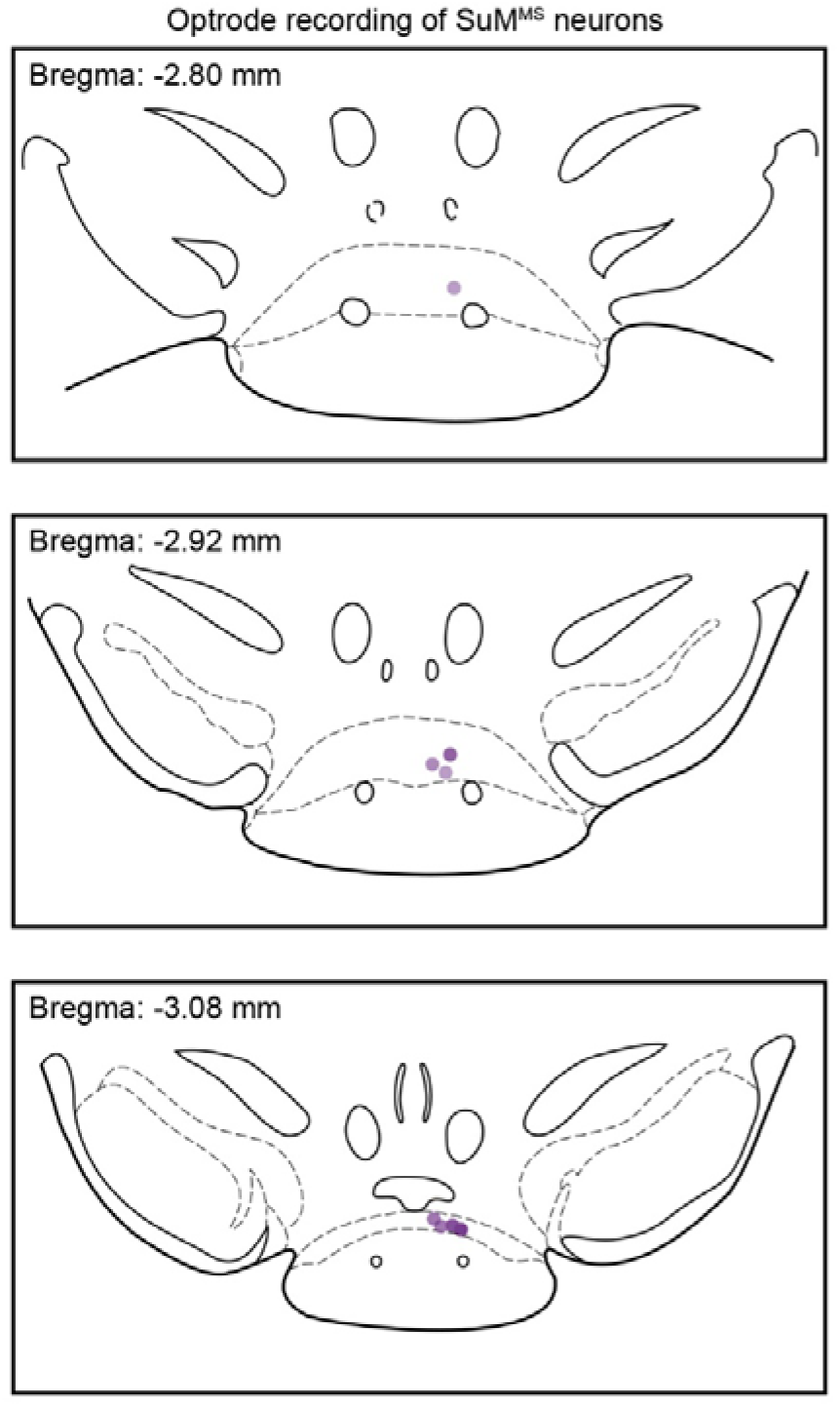
Locations of tetrodes after optrode recordings, *n* = 8 mice.

